# A Mild Keratin 1 Decrease Leads to Cell Death and Increased Oxidative Stress in B16-F10 Melanoma Cell Lines

**DOI:** 10.1101/2020.06.18.159509

**Authors:** Yujia Li, Mingchao Zhang, Weihai Ying

**Author notes:** Corresponding author: Weihai Ying, Ph.D., Professor, School of Biomedical Engineering and Med-X Research Institute, Shanghai Jiao Tong University, 1954 Huashan Road, Shanghai, 200030, P.R. China.

## Abstract

Keratins play multiple significant biological roles in epithelium. K1 / keratin 10 (K10) heterodimer is a hallmarker for keratinocyte differentiation. While keratins are absent in normal melanocyte, keratins have been found in both melanoma cell lines and human melanoma. The biological significance of the keratins in melanoma cells has remained unclear. In our current study we applied K1 siRNA to investigate the biological significance of the K1 in B16-F10 melanoma cells. We found that as low as a 16% decrease in the K1 level led to significant increases in both apoptosis and necrosis of the cells. Moreover, the mild K1 decrease led to significant increases in both dichlorofluorescein (DCF) and ethidium signals - two indicators of oxidative stress - in the cells. Collectively, our findings have provided the first evidence indicating both a critical role of the K1 in maintaining the survival of melanoma cells and an important role of the K1 in modulating the oxidative stress state of the cells. These findings have exposed new functions of keratins in cancer cells, suggesting that K1 may become a novel therapeutic target for melanoma.

## Introduction

Melanoma is the most fatal form of skin cancer [10]. Up to 4% of melanoma patients are stage IV patients with a 6% 5-year survival rate [2]. While our understanding on the pathological mechanisms of melanoma has been significantly improved during the last several decades, a significant portion of melanoma patients do not respond to systemic therapy [7]. It is of both theoretical and clinical significance to investigate novel therapeutic targets for the disease.

Keratins play multiple significant biological roles including intermediate filament formation in epithelium [9]. Keratin 1 (K1) - keratin 10 (K10) heterodimer is a hallmarker for keratinocyte differentiation [16]. K1 mutations are also associated with inherited skin diseases including epidermolytic ichthyosis, palmar-plantar keratoderma, and ichthyosis with confetti [6, 13, 14]. Several keratins, including K5, K7, K8/K18, K19, and K20, are of significance for immunohistochemical diagnosis of carcinomas [9]. It is warranted to further elucidate novel biological properties of keratins.

While keratins are absent in normal melanocytes [11], keratins have been found in both melanoma cell lines and human melanomas [3–5, 8, 15]: Several subtypes of keratins, including K1 and K10, have been found in both B16-F10-F1 melanoma cells and B16-F10-F10 melanoma cells [5]; and keratins have also been found in human melanomas [3, 4, 8, 15]. However, the biological significance of the keratins in the melanoma cells has remained unknown.

In our current study, we applied K1 siRNA to decrease the K1 in B16-F10 melanoma cell lines to determine the roles of K1 in the survival and oxidative stress state of the cells. Our study has provided the first evidence indicating a crucial role of K1 in maintaining the survival and oxidative stress state of the melanoma cells.

## Materials and Methods

### Materials

All chemicals were purchased from Sigma (St. Louis, MO, USA) except where noted.

### Cell Cultures

B16-F10 cells were plated into 24-well or 6-well culture plates in Dulbecco’s modified Eagle medium (HyClone, Logan, UT, United States) supplemented with 10% fetal bovine serum (Gibco, Carlsbad, CA, United States), 100 units/ml penicillin and 100 μg/ml streptomycin at 37 °C in a humidified incubator under 5% CO_2_.

### Western blot assays

The lysates of the cells were centrifuged at 12,000 g for 20 min at 4 °C. The protein concentrations of the samples were quantified using BCA Protein Assay Kit (Pierce Biotechonology, Rockford, IL, USA). Thirty μg of total protein was electrophoresed through a 10% SDS-polyacrylamide gel, which was then electrotransferred to 0.45 μm nitrocellulose membranes (Millipore, CA, USA). The blots were incubated with a monoclonal Anti-Cytokeratin 1 (ab185628, Abcam, Cambridge, UK) (1:4000 dilution) or β-tubulin (1:1000, Abcam, Cambridge, UK) with 0.05% BSA overnight at 4 °C, then incubated with HRP conjugated Goat Anti-Rabbit IgG (H+L) (1:4000, Jackson ImmunoResearch, PA, USA) or HRP conjugated Goat Anti-mouse IgG (1:2000, HA1006, HuaBio, Zhejiang Province, China). An ECL detection system (Thermo Scientific, Pierce, IL, USA) was used to detect the protein signals. The intensities of the bands were quantified by densitometry using Image J.

### Determinations of ROS

For Dichlorofluorescin (DCF) assay, 2,7-Dichlorofluorescin diacetate (DCFH-DA, Beyotime, China), a ROS-specific fluorescent probe, was used to measure total intracellular ROS levels. Cells were incubated with 20 μM DCFH-DA dissolved in DMEM without fetal bovine serum (FBS) for 20 min at 37°C in a dark room. After washing with PBS three times, the cells were analyzed by flow cytometry (FACSAria; Becton Dickinson, Heidelberg, Germany) to detect the mean fluorescence intensity (MFI) with an excitation wavelength of 488 nm and an emission wavelength of 525 nm.

For dihydroethidium (DHE) assay, DHE (Beyotime, China) was diluted with DMEM without FBS to the final concentration of 5 μmol/L. Cells were incubated with DHE in the incubator at 37 °C for 30 minutes. After washes with PBS, the cells were observed under a fluorescence microscope.

### Transfection of the B16-F10 melanoma cells with K1 siRNA

The cells were transfected with K1 siRNA using Lipofectamine 3000 following the manufacturer’s instructions (Thermo Fisher Scientific, Waltham, MA, USA). The sequences of the mouse keratin siRNA were CUCCCAUUUGGUUUGUAGCTT and UGACUGGUCACUCUUCAGCTT (GenePharma, Shanghai, China).

### Statistical analyses

All data are presented as mean ± SEM. Data were assessed by one-way ANOVA, followed by Student - Newman - Keuls *post hoc* test, except where noted. *P* values less than 0.05 were considered statistically significant.

## Results

We determined the effects of K1 siRNA on the K1 level of B16-F10 melanoma cells, showing that K1 siRNA led to a significant, approximately 16% decrease in the K1 level (**Figure 1**). We further determined the effects of the K1 siRNA on B16-F10 cell survival, showing that K1 siRNA treatment led to increased early-stage (Annexin V^+^/AAD^−^) and late-stage apoptosis (Annexin V^+^/AAD^+^) as well as necrosis (Annexin V^−^/AAD^+^) of the cells (**Figures 2A** **and** **2B**).

**Figure 1.**
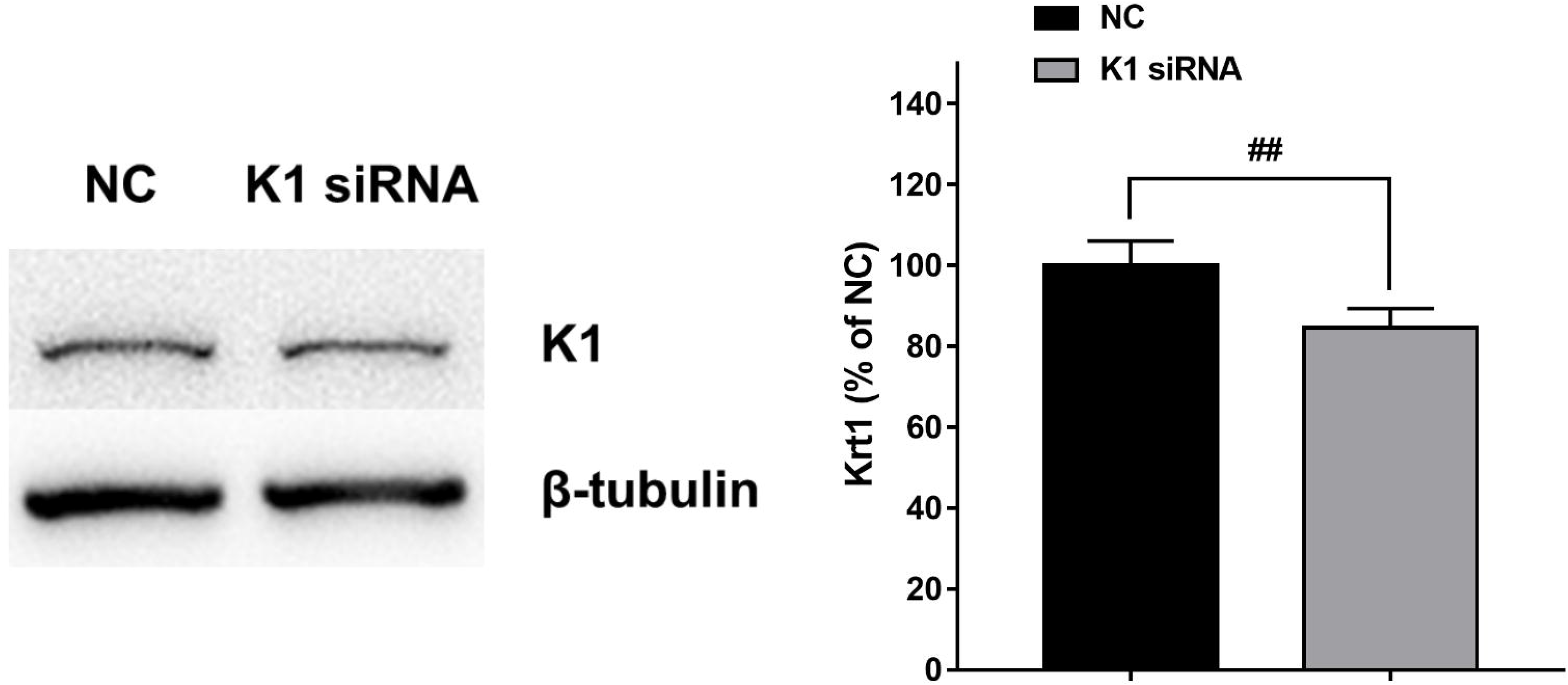
K1 siRNA treatment led to a mild decrease in theK1 level of B16-F10 melanoma cells. Twenty-four hrs after the cells were transfected with K1 siRNA, Western blot assay was conducted to determine the K1 level in the cells. ##, *P* < 0.01, N = 6.

**Figure 2.**
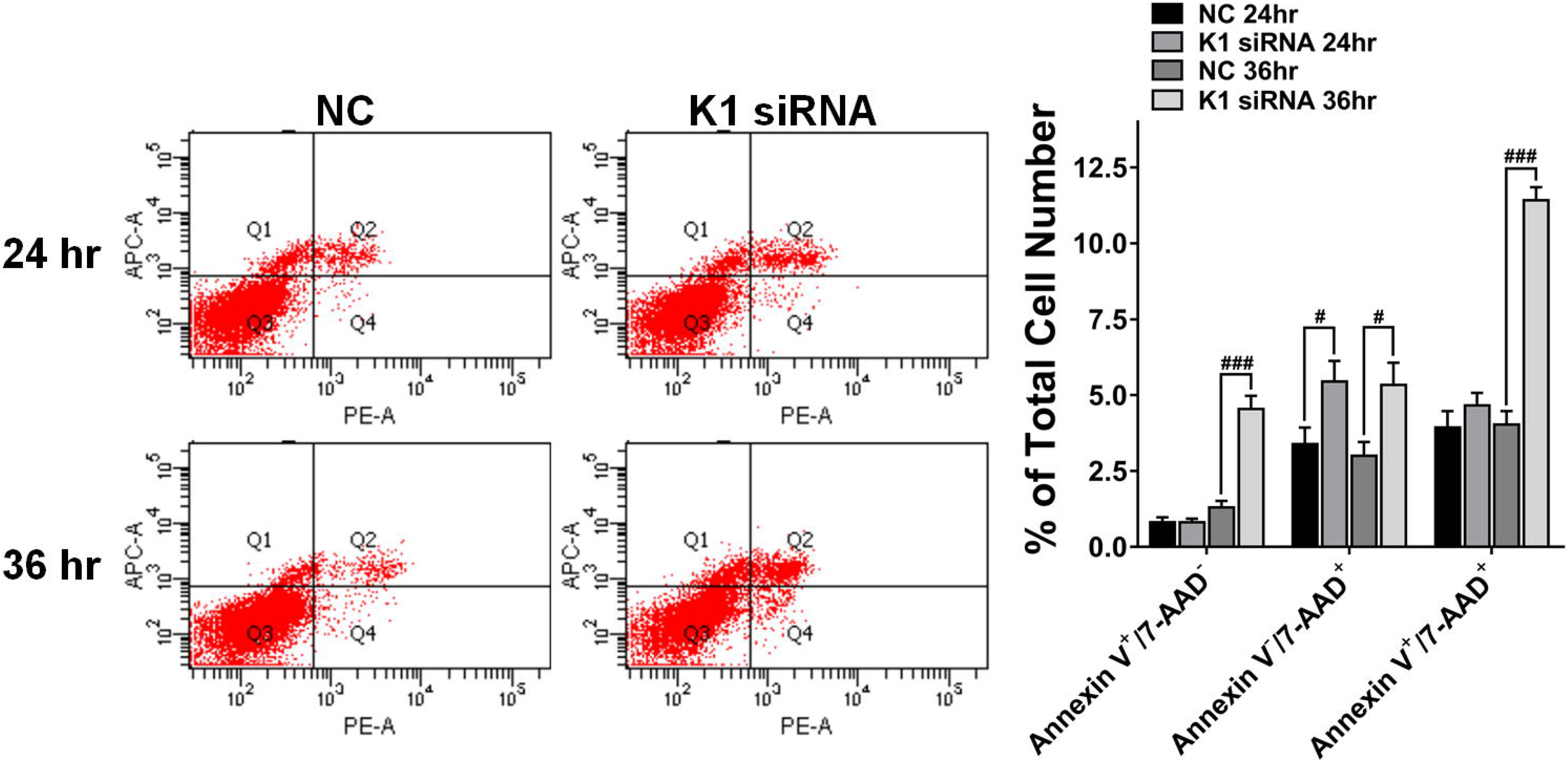
K1 siRNA treatment induced both apoptosis and necrosis of B16-F10 melanoma cells. (A) K1 siRNA treatment induced early-stage (Annexin V^+^/AAD^−^) and late-stage apoptosis (Annexin V^+^/AAD^+^) as well as necrosis (Annexin V^−^/AAD^+^) of B16-F10 cells, assessed at 36 hrs after the K1 siRNA treatment. (B) Quantifications of the effects of K1 siRNA treatment on the apoptosis and necrosis of the cells. #, *P* < 0.05; ###, *P* < 0.001. N = 9. The data were pooled from three independent experiments.

K1 siRNA induced increases in the fluorescence intensity of dichlorofluorescein (DCF) - an indicator of ROS (**Figures 3A** **and** **3B**). K1 siRNA also induced increases in the fluorescence intensity of ethidium of the cells (**Figure 4**) – an indicator of intracellular superoxide levels of the cells.

**Figure 3.**
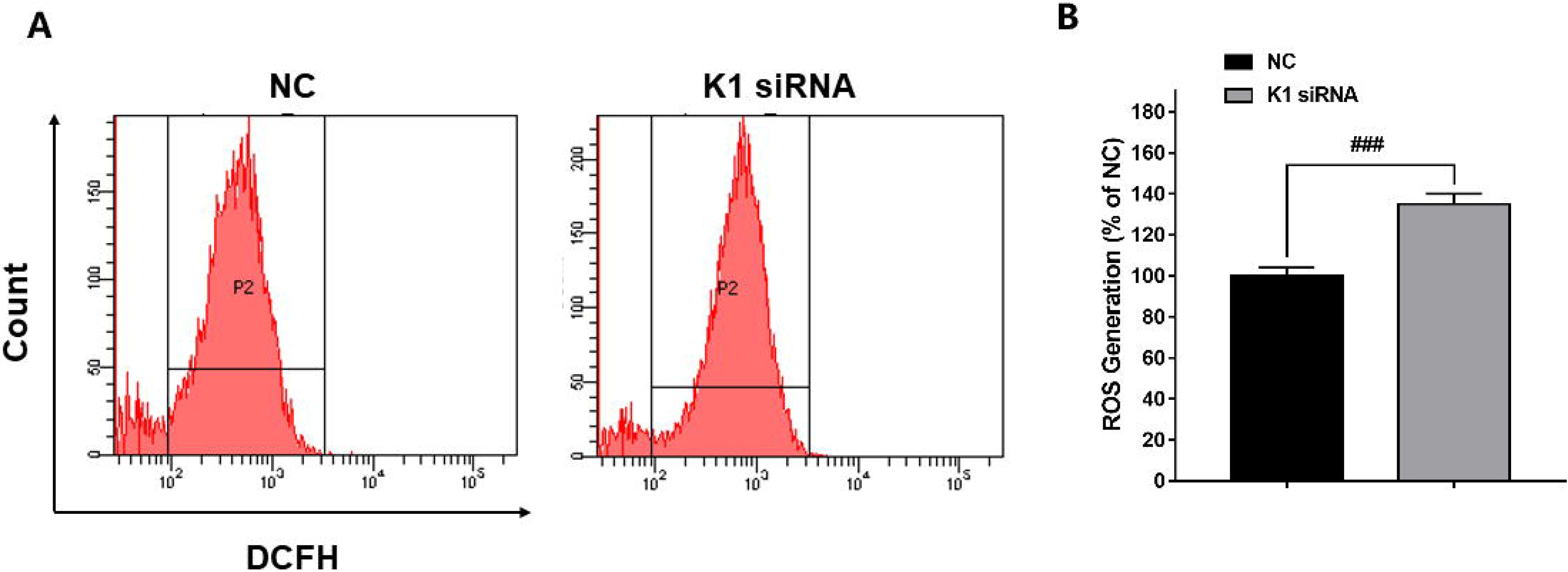
K1 siRNA treatment led to increased DCF levels of B16-F10 melanoma cells. (A) The cells treated with K1 siRNA under serum-free condition were stained with DCFH-DA. The fluorescent signals of DCF of the cells were determined by flow cytometry. (B) Quantifications of the fluorescent signals of DCF of the cells. ###, *P* < 0.001, N = 19. The data were pooled from five independent experiments.

**Figure 4.**
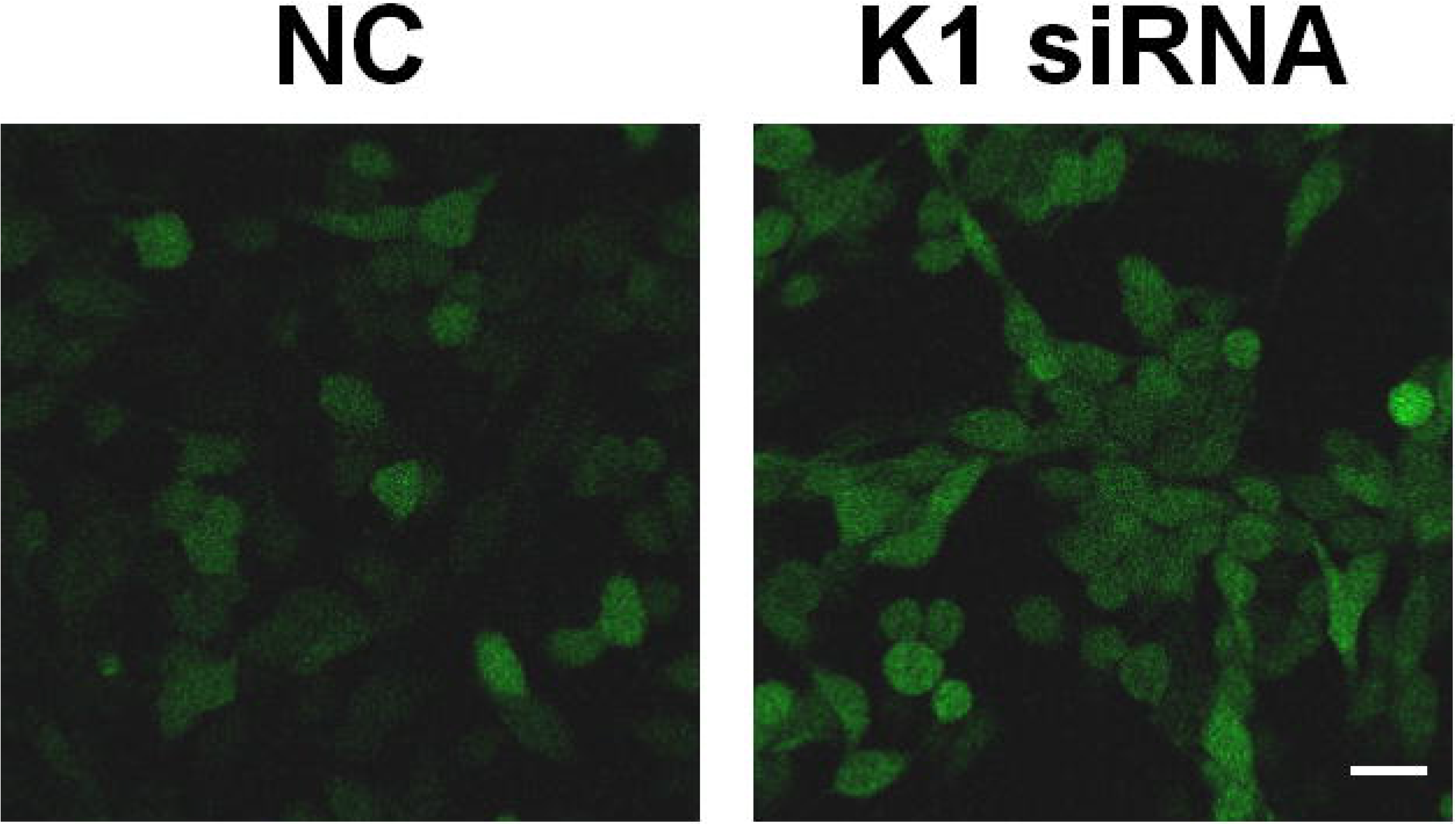
K1 siRNA treatment induced increased fluorescent signal of ethidium in B16-F10 melanoma cells. K1 siRNA treatment induced a marked increase in the fluorescent signal of ethidiumin B16-F10 cells. N = 9. The figure was a representative figure from three independent experiments.

## Discussion

The major findings of our current study include: First, a mild decrease in the K1 level of the B16-F10 melanoma cells led to both apoptosis and necrosis of the cell; and second; a mild K1 decrease of the B16-F10 melanoma cells led to a significant increase in the oxidative stress of the cell. Collectively, our study has indicated that K1 is critical for the survival of the melanoma cells.

Keratins are absent in normal melanocytes. In contrast, several subtypes of keratins, including K1 and K10, have been found in melanoma cells [5]; and several subtypes of keratins have also been found in human melanomas [3, 4, 8, 15]. However, the biological significance of the keratins in the melanoma cells has remained unknown. Our study has shown that a very mild K1 decrease is sufficient to induce significant increases in both apoptosis and necrosis of the melanoma cells. This finding has indicated significant promise of K1 as a therapeutic target for melanoma.

Oxidative stress is a key factor in cell death [12, 17]. There has been no information regarding the relationship between K1 and oxidative stress. Our study has provided first evidence indicating that K1 plays an important role in maintaining the oxidative stress state of B16-F10 melanoma cells: A mild decrease in K1 level is sufficient to induce increased oxidative stress in B16-F10 melanoma cells. This finding has also suggested a novel biological property of K1. It is warranted to further investigate the mechanisms by which K1 affects the oxidative stress of the cells. Due to the important roles of oxidative stress in cell death under numerous situations [12, 17], the mildly decreased K1 may lead to the cell death by increasing oxidative stress.

Our study has indicated the melanoma cells are highly vulnerable to the decrease in K1, suggesting that K1 may be a novel valuable target for treating melanoma cells. RNAi approach has become a promising technology for treating multiple diseases [1]. It is expected that RNAi approach is capable of producing at least a mild decrease in the K1 levels of melanoma cells, which may produce therapeutic effects on melanomas.

## Acknowledgment

The authors would like to acknowledge the financial support by two research grants from a Major Special Program Grant of Shanghai Municipality (Grant # 2017SHZDZX01) (to W.Y.) and a Major Research Grant from the Scientific Committee of Shanghai Municipality #16JC1400502 (to W.Y.).

